# The Role of the Human Brain Neuron-Glia-Synaptic Composition in Forming Resting State Functional Connectivity Networks

**DOI:** 10.1101/2021.07.01.449170

**Authors:** Sayan Kahali, Marcus E. Raichle, Dmitriy A. Yablonskiy

**Affiliations:** Department of Radiology, Washington University School of Medicine, St. Louis, MO, USA; Department of Neurology, Washington University School of Medicine, St. Louis, MO, USA

**Keywords:** functional connectivity, hierarchy of functional networks, brain cellular circuits, quantitative Gradient Recalled Echo MRI, neurons, glia cells, synapses, Human Connectome Project

## Abstract

While significant progress has been achieved in studying resting state functional networks in a healthy human brain and in a wide range of clinical conditions, many questions related to their relationship to the brain’s cellular constituents remain open. Herein we use quantitative Gradient-Recalled-Echo (qGRE) MRI for mapping human brain cellular composition, and BOLD (blood oxygen level dependent) MRI to explore how the brain cellular constituents relate to resting state functional networks. Results show that the BOLD-signal-defined synchrony of connections between cellular circuits in network-defined individual functional units is mainly associated with the regional neuronal density, while the between-functional-units connectivity strength is also influenced by the glia and synaptic components of brain tissue cellular constituents. These mechanisms lead to a rather broad distribution of resting state functional networks properties. Visual networks with the highest neuronal density (but lowest density of glial cells and synapses) exhibit the strongest coherence of BOLD signal, as well as the strongest intra-network connectivity. The Default Mode Network (DMN) is positioned near the opposite part of the spectrum with relatively low coherence of the BOLD signal but a remarkably balanced cellular content enabling DMN prominent role in the overall organization of the brain and the hierarchy of functional networks.

## INTRODUCTION

Resting state functional brain connectivity has become an established area of research in the arena of cognitive neuroscience and its related applications. Functional connectivity refers to the statistical correlation between temporally coherent low-frequency spontaneous fluctuations of the resting state functional MRI (rs-fMRI) signal in different brain regions (Biswal, Yetkin et al. 1995) and provides insights into large-scale brain circuit organization (Fox and Raichle 2007, Buckner, Krienen et al. 2013). The rs-fMRI signal is–acquired with MRI sequences sensitive to the Blood Oxygen Level Dependent (BOLD) effect (Ogawa, Lee et al. 1990) and identifies consistent resting-state networks (Damoiseaux, Rombouts et al. 2006, Yeo, Krienen et al. 2011, Van Essen, Smith et al. 2013) that play important roles in normal brain function and various neurological conditions, such for example, as normal aging (Andrews-Hanna, Snyder et al. 2007) and Alzheimer disease (Buckner, Snyder et al. 2005).

An important question in understanding the physiological basis of resting state functional connectivity is its relationship to brain structural connectivity and brain cellular composition. In the human brain, the structural connectivity issue is usually addressed by studying trajectories of brain white matter (WM) tracts with Diffusion Tensor Imaging (DTI) tractography (Toosy, Ciccarelli et al. 2004, De Luca, Beckmann et al. 2006, Greicius, Supekar et al. 2009, Uddin 2013). However, the direct structural connectivity through WM is not the sole mechanism underlying functional connectivity (Honey, Sporns et al. 2009) (Ding, Huang et al. 2018).

While DTI is sensitive to structural connections governed by WM tracts, it does not usually have enough sensitivity to resolve existing neuronal connections through the brain cortical regions, which are characterized by a complex network of interconnected and intersected neuronal processes (axons and dendrites). It is also understood that functional connectivity of the brain is not uniquely dependent on the neurons, it also relies on glial cells which significantly influence structural and functional connectivity (Fields, Woo et al. 2015), providing metabolic and regulatory support for neurons (Ullian, Sapperstein et al. 2001, Pannasch, Vargova et al. 2011) (Pellerin and Magistretti 2012). Astrocytes are responsible for increasing the amount of mature and functional synapses (Ullian, Sapperstein et al. 2001, Pannasch, Vargova et al. 2011). Moreover, neuron-glia cross-talk leads to synaptic formation and remodeling (Stogsdill and Eroglu 2017). It has also been observed that neurons generate weak synapses in the absence of glia (Araque and Navarrete 2010, Pannasch, Vargova et al. 2011). Hence, incorporating information on cellular composition into brain structural connectivity can provide crucial information for understanding the brain functional connectivity network formation and functioning.

In this paper, to identify brain cellular structure we use data obtained by means of quantitative Gradient Recalled Echo (qGRE) MRI technique (Ulrich and Yablonskiy 2016). The qGRE method is based on the gradient recalled echo MRI with multiple gradient echoes and data analysis allowing separation of tissue cellular specific (R2t*) GRE signal relaxation from relaxation caused by the *baseline* BOLD (blood oxygen level dependent) mechanism (Ogawa, Lee et al. 1990, Yablonskiy and Haacke 1994) and adverse effects of magnetic field inhomogeneities (Yablonskiy 1998) corrected using a VSF (voxel spread function) method (Yablonskiy, Sukstanskii et al. 2013). We also use a quantitative relationships established in Wen et al. (Wen, Goyal et al. 2018) between brain cellular composition and the R2t* metric of the qGRE MRI signal. Based on the analysis of the genetic information from the Allen Human Brain Atlas, Wen et al. (Wen, Goyal et al. 2018) identified several networks of gene expression profiles coherently expressed across brain anatomical structures and established their association with the R2t* metric of the qGRE signal in these structures. Data showed the strongest association between R2t* and genes that are associated with ion channels primarily distributed along neuronal processes, including axons (myelinated and not myelinated) and dendrites which typically occupy over 85% of the space comprised by neurons. Noting these findings, it was demonstrated (Wen, Goyal et al. 2018) that the in vivo R2t* measurement largely reflects a transcriptional correlate for major parts of the neuronal cell bodies and processes, thus representing the neuronal contribution to brain tissue cellular composition. Moreover, a quantitative relationship between the R2t* metric of the qGRE signal in a healthy human brain cortical Grey Matter (GM) and a neuronal density index *Y_neuron_* that serves as a proxy for the brain tissue neuronal density was established in (Wen, Goyal et al. 2018):

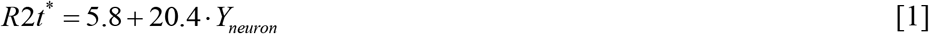

where R2t* is measured in sec^-1^ and the neuronal density index Y_neuron_ is a dimensionless parameter (0 < Y_neuron_ < 1) proportional to the tissue neuronal density. This relationship is illustrated in Appendix Figure A1. Further analysis (Wen, Goyal et al. 2018) also established relationships in healthy human brain cortical GM between *Y_neuron_* and indices characterizing densities of glia cells (*Y_glia_*) and synapses (*Y_synapse_*):

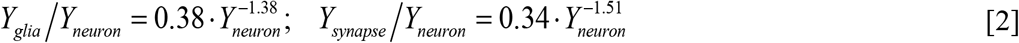

thus, corroborating previous histology-based reports (Cullen, Gilroy et al. 2010, Herculano-Houzel 2014) that healthy human brain regions with higher neuronal density have relatively low densities of glia cells and synapses. Herein we combine qGRE-derived brain cellular information with resting state BOLD data from the Human Connectome Project (HCP) to establish a quantitative relationship between resting state functional connectivity and human brain cortical cellular composition. We show that while the synchrony of the brain cellular circuits in network-defined functional units is mainly associated with the regional neuronal density, the strength of the inter-unit within-network functional connectivity has significant association not only with units’ neuronal content but also with their glia and synaptic constituents. These cellular-functional associations are most prominent in the infra-slow frequency range (0.01 – 0.16 Hz) of brain activity as determined by the BOLD signal.

## METHODS

The functional connectivity data were obtained from the HCP1200 dataset (February, 2017 release). The qGRE data were obtained from healthy control participants, with local IRB approval from Washington University in Saint Louis. Written informed consent was obtained from all participants. All methods were performed in accordance with relevant guidelines and regulations.

### Resting State Functional Connectivity Data

The HCP1200 dataset was recorded on young adults aged between 22-35 (Van Essen, Smith et al. 2013). In this study, we used 183 subjects (about 20% of total available) that had 4, 15-minute RS-fMRI scans. The fMRI scans were preprocessed using HCP structural (PreFreeSurfer, FreeSurfer, and PostFreeSurfer) and functional (fMRIVolume and fMRISurface) pipelines. The ICA+FIX pipeline was applied to refine the rs-fMRI data to regress out the spatially specific temporal artefacts, i.e., scanner artefacts, subject movement, breathing and cardiac pulsation. The fMRI data were then summarized in 91,282 grayordinates which are cortical grey matter surface vertices and subcortical grey matter voxels(Glasser, Sotiropoulos et al. 2013). The data in each grayordinate obtained from 4 scans of each subject (1200 timepoints ×4 runs) were concatenated after performing de-meaning and variance normalization. The concatenated time series data for each subject were grouped into 300 ROIs (i.e., functional units) using Schaefer’s et al. (Schaefer, Kong et al. 2018) local-global parcellation of the cortical grey matter identified by employing the gradient-weighted Markov Random Field (gwMRF) model which fuses local gradient and global similarity approaches. Signals from these time-series data for all vertices in each ROI were averaged together and each ROI for further analysis was represented by a single signal (1200 timepoints ×4 runs). These time-series data for each of 183 subjects were concatenated together for time domain network analysis (1200 timepoints ×4 runs × 183 subjects).

To quantitatively characterize rs-fMRI signal (i.e. the statistical correlation between temporally coherent spontaneous fluctuations of the rs-fMRI signal in different brain regions), we introduce a short-range signal coherence (*signal coherence in individual functional units*) and a long-range signal coherence (*signal coherence between functional units*).

To represent the signal coherence of the resting state functional *BOLD signal from each individual functional unit* (*designated below as Region Of Interest, i.e. ROI*) we calculated the Standard Deviation (*STD_i_*) of the average signals combined from individual vertices in each functional unit *i* (*i* =1, 2,…, 300). Due to normalization, the *STD* of each signal from the vertices is equal to one, such that the *STD_i_* of the combined signal represents *the coherence* of the individual signals in the ROI (*STD* = 1 if signals from vertices are coherent, and *STD* = 0 in there is no correlation between them). Then we represent the BOLD signal coherence for each network as a mean value of *STD_i_* of all *ROI_i_* belonging to a given network:

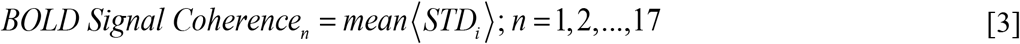

The resting state functional Connectivity Strength (CS*_i,j_*) between two ROIs (*i* and *j*), i.e. *the signal coherence between functional units*, is calculated as a Pearson Correlation Coefficient between two time series data from *ROI_i_* and *ROI_j_*, each having 1200 timepoints × 4 runs x 183 subjects.

To characterize the intra-network connectivity, we introduce the *intra-Network Connectivity Strength NCS*_n_ by calculated weighted average *CS_i,j_* between all ROIs belonging to a given network:

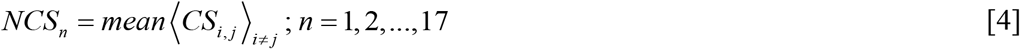

Since ROIs contain different number of vertices, all ROIs in this calculation are weighted by the number of corresponding vertices.

The frequency content of rs-fMRI signal was analyzed by converting the time-series data to frequency domain by performing a Fourier transformation on time-series data from each ROI independently, for each session per subject. The Fourier transformed data were then averaged across all sessions and all subjects, before separation into preselected frequency bins with the widths of 0.014 Hz and 0.0047 Hz. Signal from each bin was then converted back to the time domain using inverse Fourier transform.

### Quantitative Gradient Recalled Echo (qGRE) MRI Data Analysis

To identify brain cellular structure we used data obtained by means of quantitative Gradient Recalled Echo (qGRE) MRI technique (Ulrich and Yablonskiy 2016). The qGRE method is based on the gradient recalled echo MRI with multiple gradient echoes and data analysis allowing separation of tissue cellular specific (R2t*) GRE signal relaxation, from relaxation caused by the baseline Blood Oxygen Level Dependent (BOLD) mechanism and adverse effects of magnetic field inhomogeneities (Yablonskiy 1998). We used MRI data obtained from 16 healthy volunteers aged between 23-35 from a previously published study (Zhao, Wen et al. 2016). These volunteers were not a part of the HCP sample, but their age range was selected to match HCP resting state functional MRI data. In Zhao et al., (Zhao, Wen et al. 2016) MRI image data were obtained using a 3T Trio MRI scanner (Siemens, Erlangen, Germany) using a 32-channel phased-array RF head coil. Data acquisition was done by means of 3D Gradient Recalled Echo (GRE) MRI sequence with 10 gradient echoes, followed by a navigator for correcting physiological fluctuations (Wen, Cross et al. 2015). The sequence parameters were flip angle (FA) = 30°, repetition time (TR) = 50 ms, first echo time (TE_1_) = 4 ms, echo spacing ΔTE = 4 ms, voxel size of 1 × 1 × 2 mm^3^ and acquisition time of 11 minutes. Field inhomogeneity effects were corrected using a Voxel Spread Function (VSF) method (Yablonskiy, Sukstanskii et al. 2013).

Data analysis was performed with a stand-alone computer with in-house developed programs written in MatLab (MathWorks Inc., Natick, MA, USA). After phase correction, k-space data from each radio frequency (RF) channel were converted to the spatial domain, and the 3D spatial Hanning filter was applied to reduce Gibbs ringing artifacts and signal noise. To achieve optimal model parameter estimations, the multi-channel data (ch = 1, 2,…, M) were combined according to the following algorithm allowing the most accurate model parameters evaluation (Quirk, Sukstanskii et al. 2009, Luo, Jagadeesan et al. 2012):

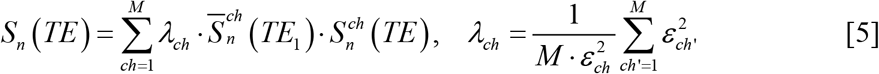

Where 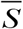 denotes the complex conjugate of S; index n represents the voxel position in space; λ_ch_ are weighting factors, and ε_ch_ are noise amplitudes (r.m.s.).

A theoretical model of BOLD (blood oxygen level dependent) contrast (Yablonskiy and Haacke 1994) was used to differentiate the contribution of tissue-cellular-specific relaxation (R2t*) and BOLD contributions to the total R2* relaxation:

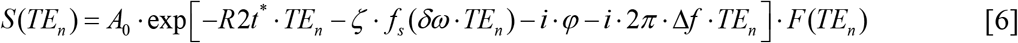

where *A*_0_ is the signal amplitude; *δω* is the characteristic frequency determined by the susceptibility difference between deoxygenated blood and surrounding tissue; ζ is the volume fraction of deoxygenated blood; nonlinear function *f_s_* (*δω*·*TE*) accounts for the BOLD effect (Yablonskiy and Haacke 1994); *φ* and Δ*f* are phase and frequency shifts (dependent on tissue structure and also macroscopic magnetic field created mostly by tissue/air interfaces), and the function *F*(*TE*_n_) describes the effect of macroscopic magnetic field inhomogeneities (Yablonskiy 1998). Herein, *F*(*TE*_n_) was calculated by a voxel spread function (VSF) method (Yablonskiy, Sukstanskii et al. 2013) and we used a mathematical expression for the function *f_s_* in terms of a generalized hypergeometric function (Yablonskiy, Sukstanskii et al. 2013) _1_*F*_2_:

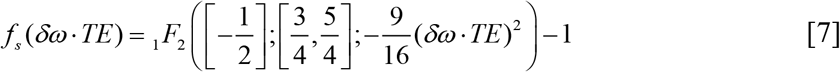

### qGRE-Defined Brain Cellular Structure and Cellular Associations

For our analysis, the R2t*-based calculated cellular indices for each subject, Eqs. [1] and [2], were projected on the cortical surface through the surface-based registration using the grayordinates. The grayordinates were then combined into 300 ROIs based on the Schaefer’s et al. parcellation (Schaefer, Kong et al. 2018). After that, the results for all 16 subjects were averaged for each of the ROIs producing 300 sets of cellular indices (*Y_neuron_*, *Y_gila_* and *Y_synapse_*).

To establish a joint contribution of two distinct ROIs (*i* and *j*) to their resting state functional Connectivity Strength CS*_i,j_*, we consider six *Cellular Association* (*CA_i,j_*) matrices depending on their cellular composition such as, neuron-neuron, glia-glia, synapse-synapse, neuron-glia, neuron-synapse, and glia-synapse which can be defined on 300 ROIs as follows:

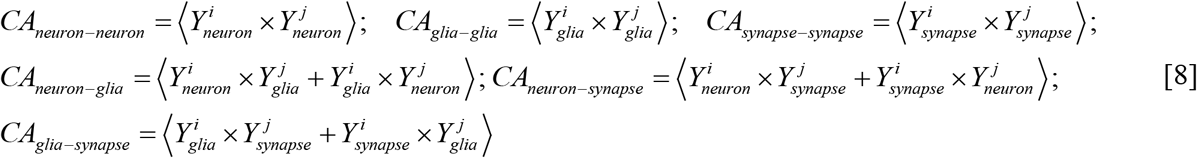

For each network we also introduce *internal (intra-network) Cellular Association Indices* for six types of associations: neuron-neuron, glia-glia, synapse-synapse, neuron-glia, neuron synapse and glia-synapse, whereas each of these indices is calculated as a *weighted average of CA_i,j_* (*i* ≠*j*) in Eq. [8] across all combinations of ROIs in the *n*-th network (similar to *NCS* in Eq. [4]). All ROIs in these calculations are weighted by their number of voxels.

### Analysis based on T1w/T2w approach

To further understand the role that different parts of the neuron play in formation of brain functional connectivity, we can compare our R2t*-based results with a ratio of T1-weighted (T1w) to T2-weighted (T2w) images as a proxy related to cortical tissue myelin content proposed by Glasser and Van Essen (Glasser and Van Essen 2011). While this proxy does not represent a quantitative measure of myelin content, for example as discussed in (Lazari and Lipp 2021), it was successfully used for mapping cytoarchitecture of human cortical areas (Glasser, Coalson et al. 2016). By using T1w and T2w data from the same HCP subjects that are used in this paper for rs-fMRI analysis, we have calculated a Myelin Index (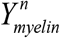; *n* = 1,2,…,17, as a ratio of T1w to T2w images) for all ROIs in Yeo’s networks. Similar to cellular association indices, we have also introduced indices characterizing myelin association in each network.

## RESULTS

### The Strength of Resting State Functional Networks is Significantly Associated with the Neuron-Neuron, Neuron-Glia and Neuron-Synaptic Structural Circuits in the Human Brain Cortex

In this paper we provide data analysis based on a structure of 300 gwMRF ROIs combined in 17 resting-state networks developed by Yeo et al (Yeo, Krienen et al. 2011, Schaefer, Kong et al. 2018). These networks are presented in Figure 1. We have selected a gwMRF brain parcellation scheme (Schaefer, Kong et al. 2018) because it exhibits an improved functional connectivity homogeneity compared with other parcellations (Craddock, James et al. 2012, Shen, Tokoglu et al. 2013, Glasser, Coalson et al. 2016, Gordon, Laumann et al. 2016) and provides a sufficient number of parcels (ROIs) to delineate brain cortical anatomical structures as discussed by Van Essen et al. (Van Essen, Glasser et al. 2012).

**Figure 1.**
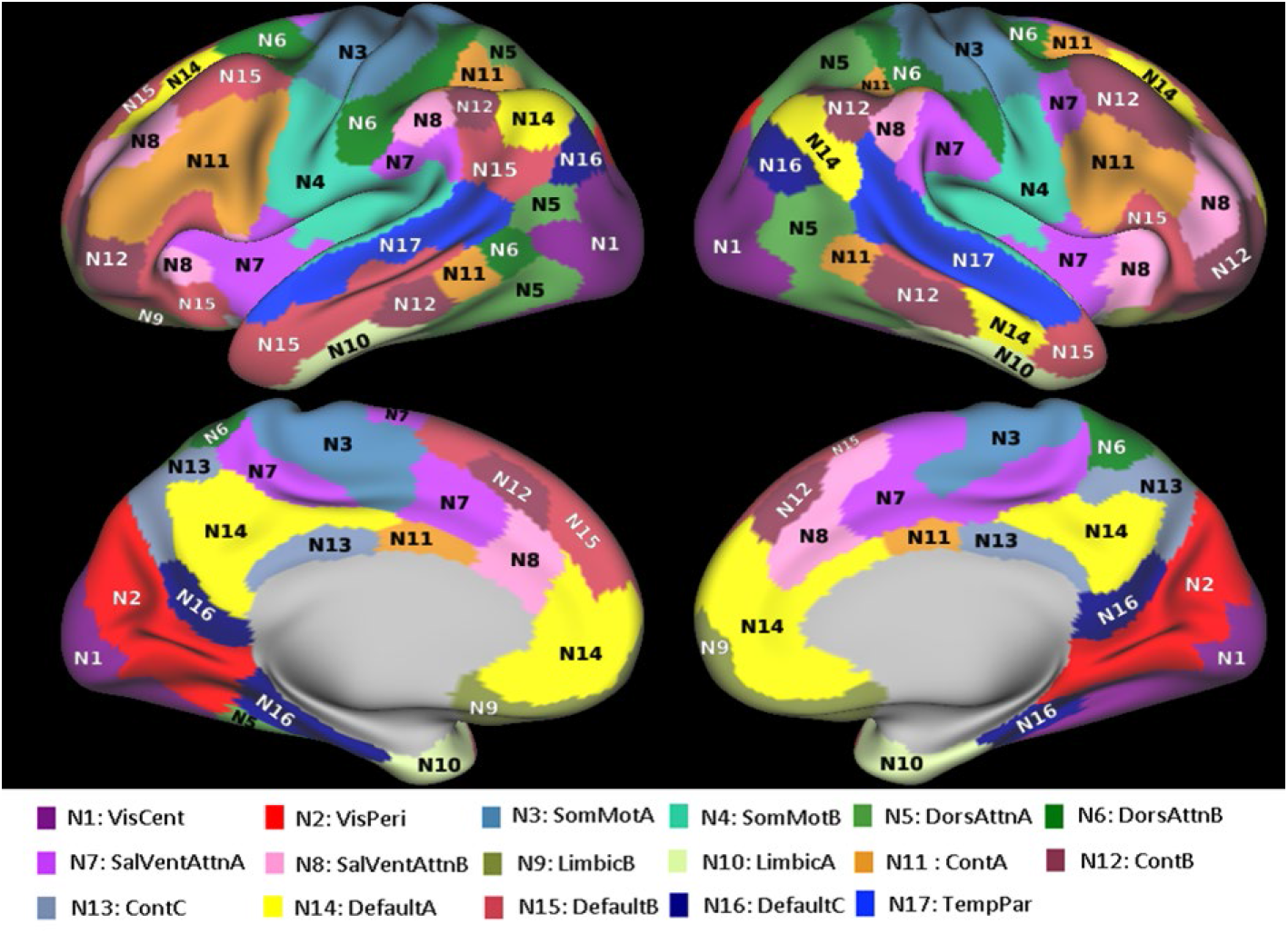
Network parcellation of Yeo’s 17-networks. The 17-networks include the following regions: N1: VisCent - Visual A (22 ROIs), N2: VisPeri - Visual B (18 ROIs), N3: SomMotA - Somatomotor A (30 ROIs), N4: SomMotB - Somatomotor B (21 ROIs), N5: DorsAttnA - Dorsal Attention A (16 ROIs), N6: DorsAttnB - Dorsal Attention B (16 ROIs), N7: SalVentAttnA - Salience/Ventral Attention A (25 ROIs), N8: SalVentAttnB - Salience/ Ventral Attention B (14 ROIs), N9: LimbicB - Limbic B (9 ROIs), N10: LimbicA - Limbic A (11 ROIs), N11: ContA - Control A (22 ROIs), N12: ContB - Control B (18 ROIs), N13: ContC - Control C (8 ROIs), N14: DefaultA - Default A (23 ROIs), N15: DefaultB - Default B (25 ROIs), N16: DefaultC - Default C (10 ROIs), N17: TempPar - Temporal Parietal (12 ROIs). Further details on networks’ structure can be found in (Yeo, Krienen et al. 2011, Schaefer, Kong et al. 2018).

Using qGRE data we have analyzed the contribution of magnetic field inhomogeneities in signal formation of all 17 networks and found that the two limbic networks are affected the most. In fact, the signals from the networks Limbic A and Limbic B had more than 88% and 80% of voxels significantly affected by the background magnetic field inhomogeneities, respectively, while all other networks had on average only 9% of “bad” voxels (see Appendix Figure A2 for details). Consequently, we omitted limbic networks from further consideration.

The theoretical approach described in the Methods section allows – based on Eqs. [1] and [2] – the calculation of neuronal, synaptic, and glia cells indices (qGRE proxy parameters proportional to the neuronal, synaptic, and glia cells densities) for each network. The R2t* maps were obtained from 16 healthy volunteers aged 23 to 35 as described in the Methods section, projected on 300 selected ROIs and averaged together. **Table 1** in the Appendix shows variation of R2t* measurements across the subjects. Images representing mean neuronal, glia and synaptic density indices are presented in **Figure 2**.

**Table 1.** Summary of R2t* and cellular densities in 15 networks. Columns 2 and 3 represent Mean and Standard deviation of R2t* distribution across 16 subjects (values are in 1/sec). Columns 4-6 represent mean values of neuronal, glia and synaptic density indices (qGRE proxy for the neuronal (NDI), synaptic (SDI) and glia (GDI) cells densities). Data in columns 4-6 correspond to the maps in **Figure 2** of the main text.

| NETWORKS | R2t* Mean | R2t* STD | NDI | GDI | SDI |
| --- | --- | --- | --- | --- | --- |
| Vis-Cent | 19.67 | 0.8 | 0.66 | 0.44 | 0.42 |
| Vis-Peri | 18.86 | 0.48 | 0.65 | 0.45 | 0.43 |
| SomMot-A | 16.95 | 0.82 | 0.58 | 0.47 | 0.45 |
| SomMot-B | 18.02 | 0.78 | 0.6 | 0.46 | 0.44 |
| DorsAttnA | 17.88 | 0.67 | 0.61 | 0.46 | 0.44 |
| DorsAttnB | 17.13 | 0.79 | 0.57 | 0.47 | 0.45 |
| SalVentAttn-A | 16.61 | 0.68 | 0.53 | 0.49 | 0.47 |
| SalVentAttn-B | 16.34 | 0.71 | 0.52 | 0.49 | 0.47 |
| Cont-A | 17.69 | 0.73 | 0.59 | 0.47 | 0.45 |
| Cont-B | 16.55 | 0.74 | 0.53 | 0.49 | 0.47 |
| Cont-C | 17.87 | 0.96 | 0.6 | 0.46 | 0.44 |
| Default-A | 16.6 | 0.71 | 0.55 | 0.48 | 0.47 |
| Default-B | 16.29 | 0.83 | 0.5 | 0.5 | 0.49 |
| Default-C | 18.06 | 0.71 | 0.6 | 0.49 | 0.44 |
| TempPar | 17.81 | 0.68 | 0.59 | 0.47 | 0.45 |

**Figure 2.**
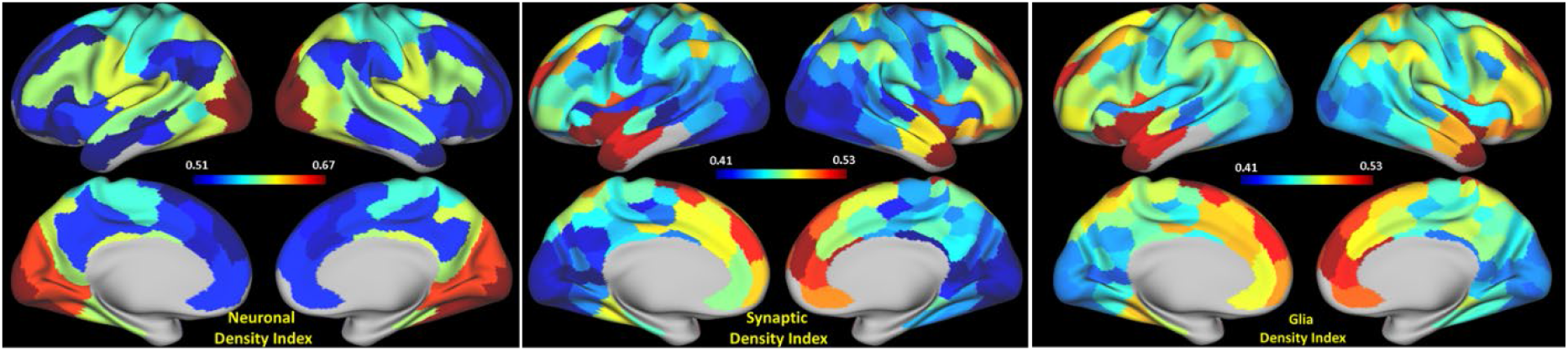
Surface maps of neuronal, glia and synaptic density indices (qGRE proxy for the neuronal, synaptic and glia cells densities) for 15 networks. Data representing the results in this figure are shown in the Appendix **Table 1**.

Plots in **Figure 3** represent correlation between BOLD signal coherence in 15 networks and neural, glia, and synaptic density indices in these networks. Data show strong positive association with mean neuronal density index and negative association with glia and synaptic density indices. *This means that neurons*, *not glia cells and synapses*, *are mostly responsible for the coherence of the BOLD signal*. The strong negative association of BOLD signal coherence with glia and synaptic density indices is due to the fact that the densities of glial cells and synapses are negatively associated with neuronal density in the corresponding regions as presented in Eq. [2].

**Figure 3.**
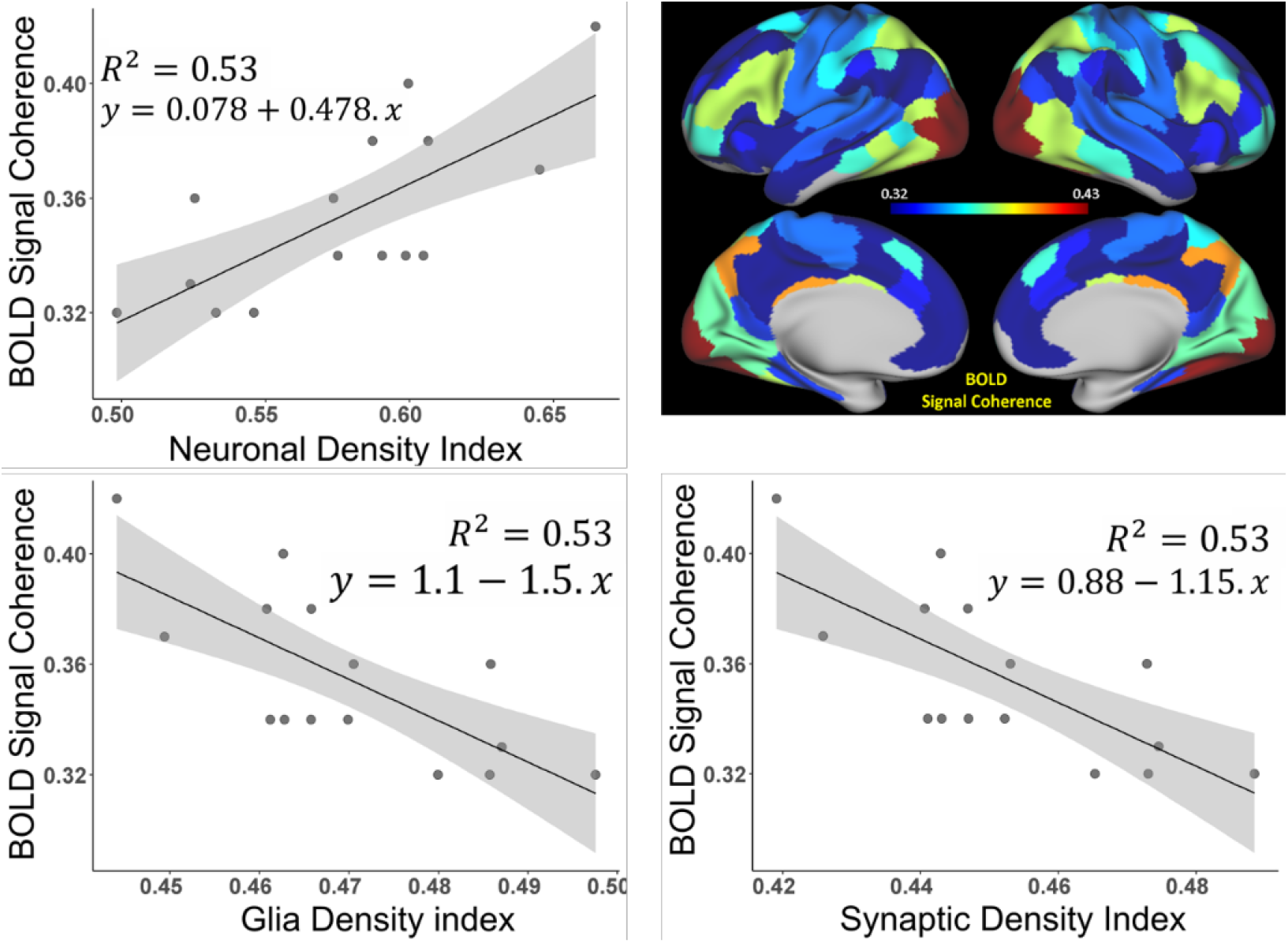
Association between BOLD signal coherence in 15 networks and neural, glia, and synaptic density indices in these networks (limbic networks are omitted). Neuronal (Y_neuron_), glia (Y_glia_), and synaptic (Y_synapse_) density indices were calculated from R2t* values using Eqs.[1] and [2] and then averaged for each network. Shaded areas – 95% confidence intervals. The R^2^ values for all three correlations are practically identical because of previously established correlated spatial distribution of neurons, glia, and synapses in the healthy human cortex (Cullen, Gilroy et al. 2010, Herculano-Houzel 2014) which is reflected by the relationship between glia (Y_glia_) and synaptic (Y_synapse_) indices to the neuronal index (Y_neuron_) by means of Eq. [2].

Even though the strength of the BOLD signal coherence in functional networks is mostly associated with the networks’ neuronal content (**Figure 3**), the functional connectivity of the brain is not solely dependent on the neurons, it also includes glial cells (Astrocyte, Microglia, Oligodendrocytes) which significantly influence structural and functional connectivity governed by neurons (Fields, Woo et al. 2015). Obviously, the factors defining the strength of functional connectivity between two ROIs are the actual neuronal pathways connecting these ROIs and the cellular composition of these ROIs. Herein we focus on the latter – the ROIs’ cellular composition.

**Figure 4** show correlations between the *intra-Network Connectivity Strength*, *NCS_n_*, Eqs. [8] and [9] and the cellular association strength *CAS^n^* of these networks for six types of cellular associations (neuron-neuron, glia-glia, synapse-synapse, neuron-glia, neuron-synapse, and glia-synapse). Data illustrates that the neuron-neuron (***CAS**_neuron-neuron_*) neuron-glia (***CAS**_neuron-glia_*) and neuron-synapse (***CAS**_neuron-synapse_*) associations show strong positive correlations suggesting these associations as dominant features contributing to the inter-ROIs resting state functional connectivity.

**Figure 4.**
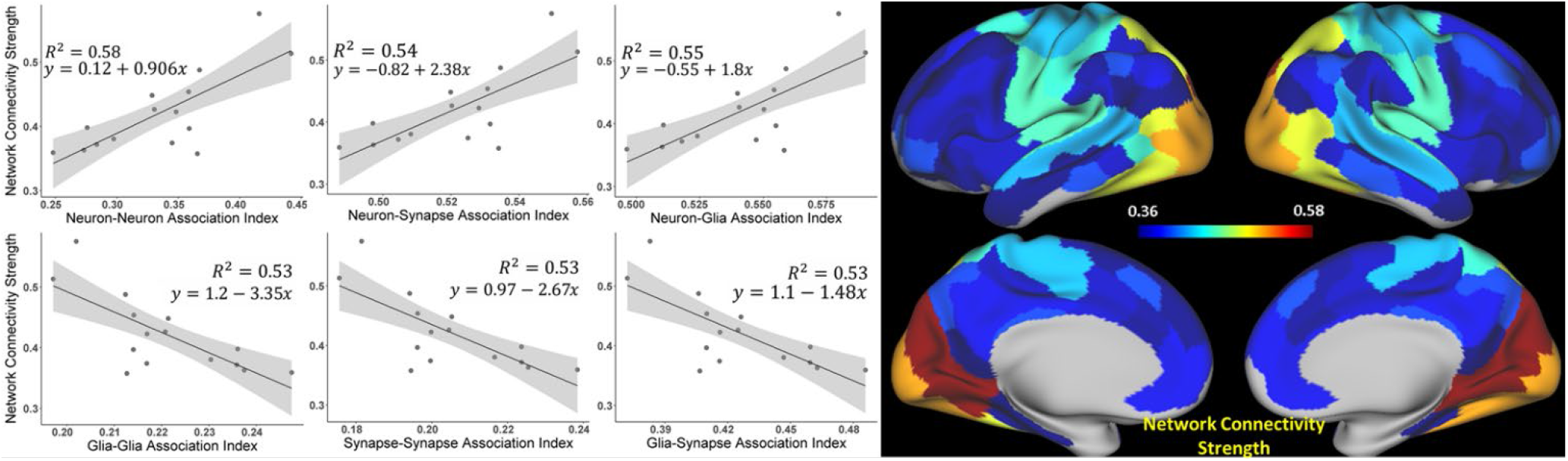
Relationships between Network Connectivity Strength (NCS) and Indices characterizing Cellular Associations between neurons, synapses, and glia cells. Plots show correlations between resting state network connectivity strength NCS_n_ and cellular associations strength *CAS^n^* (index ^n^ is omitted in the figure) of these networks for six types of cellular associations (neuron-neuron, glia-glia, synapse-synapse, neuron-glia, neuron-synapse, and glia-synapse) for 15-networks. Shaded areas – 95% confidence intervals. Images represent surface maps of NCS for 15 networks.

While data in **Figure 3** and **Figure 4** show network-wide associations between the BOLD signal coherence vs. neural density index and network connectivity strength vs. cellular association strength between cells, respectively, **Figure A3** in the Appendix shows similar associations based on 280 individual ROIs (limbic-network-associated ROIs were excluded), which are highly significant (p= E-11 and E-98 correspondingly).

We have also tested for the randomness of our correlation result in **Figure 3** (BOLD signal coherence vs. Neuronal Density Index) and **Figure 4** (Functional Connectivity (FC) vs. cellular association strength between neurons) by means of permutation test (randomization test). The permutation test for BOLD signal coherence vs. Neuronal Density Index was done by randomly permuting BOLD signal coherence 10000 times and calculating correlation with Neuronal Density Index for each permutation. The curve in **Figure A4 (b)** in the Appendix illustrates the distribution of obtained *r*-values validating a significant difference with p-value < 0.000001). Similarly, the functional connectivity strengths between 280 ROIs (excluding Limbic networks) were randomly permuted 20000 times and correlated with the cellular association strength between neurons for each permutation. The curve in the **Figure A4 (a)** in the Appendix shows the distribution of obtained *r*-values, thus validating a significant difference (p-value < 0.000001).

To demonstrate statistical sufficiency of our sample size (N=183) we ran analyses similar to those presented in **Figure 3** and **Figure 4**, using a smaller sample size (N = 100). Results presented in Appendix **Figure A5** show practically the same correlations, with only slightly smaller R^2^ values.

Data in Figure **5** show associations between cortical tissue Myelin Index calculated based on the ratio of T1w/T2w images and qGRE-based calculated Neuronal Density Index (**Figure 5a**), BOLD signal Coherence (**Figure 5b**), and Network Connectivity Strength (**Figure 5c**).

**Figure 5.**
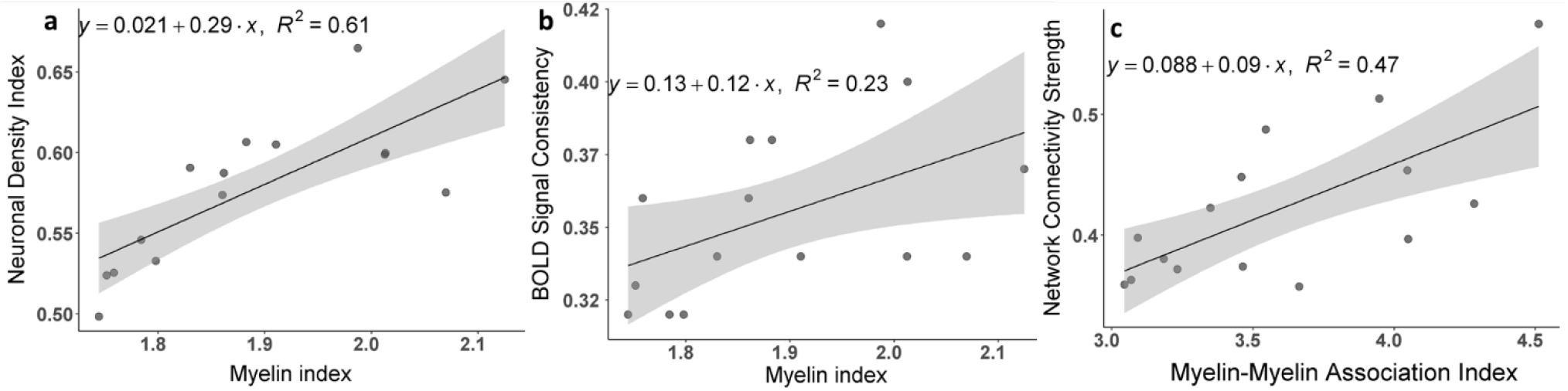
Association between qGRE and T1w/T2w –results based on 15 networks. **(a)** Correlation between Neuronal Density Index and Myelin Index. Points represent average values for individual networks. Shaded areas – 95% confidence intervals. Myelin Index was calculated based on T1w/T2w images. **(b)** Correlations between the strength of the BOLD signal coherence and average Myelin Index in the networks. **(c)** Correlation between resting state intra-network connectivity strength, NCS and Myelin-Myelin Association Index calculated based on the myelin indices in the connected ROIs (similar to that described by Eqs. [4], [8]).

### Brain cortical cellular composition shows the strongest association with the resting state BOLD signal coherence and network connectivity in the infra-slow frequency range of neuronal activity

Data in **Figure 3** show that the BOLD signal coherence varies significantly among networks and strongly correlates with the networks’ neuronal density. Similar behavior is seen for network connectivity strength which also depends on the brain cellular composition, though in a different manner; the intra-network connectivity strength shows the strongest correlation with intra-network neuron-neuron, neuron-synaptic, and neuron-glia associations between network’s ROIs. These results were obtained by analyzing the time courses of rs-fMRI signal without accounting for frequency content of this signal. At the same time, only the infra-slow (below 0.1 Hz) frequency fluctuations of the resting state BOLD signal are usually considered as a signature of the neuronal activity (Fox and Raichle 2007, Zhang and Raichle 2010, Han, Wang et al. 2011, Palva and Palva 2012, Mitra, Kraft et al. 2018, Raut, Snyder et al. 2020). To investigate the detailed relationship between cellular composition and frequency components of the resting state BOLD signal, we converted the time series data into 50 consecutive frequency bands covering frequency domain 0.01-0.60 Hz (see methods section) and analyzed different frequency components independently. The results for 15 networks are presented in **Figure 6**. **Figure 6a** shows the BOLD signal coherence (mean standard deviation of the rs-fMRI signal) in each network as a function of the rs-fMRI signal frequency. We found that while different networks exhibit different signal strength coherence, the general trend is quite similar – the strongest coherence of BOLD signal fluctuations is present in the frequencies below 0.16 Hz, with the peak between 0.01-0.03 Hz. The NCS calculated in terms of the mean intra-network correlation coefficient for each of the 15 networks as a function of rs-fMRI signal frequency content is seen in **Figure 6b,** which also displays increased connectivity strengths in a similar frequency range below 0.16 Hz, with the peak between 0.01-0.03 Hz. **Figure 6c** shows correlation between BOLD signal coherence in 15 networks and the tissue neuronal content of these networks (as in **Figure 3**) as a function of the rs-fMRI signal frequency. Interestingly, the strongest correlation exists not only in the frequency range where the BOLD signal coherence and NCS are the strongest, but in a rather broad infra-slow frequency range between 0.01 – 0.16 Hz.

**Figure 6.**
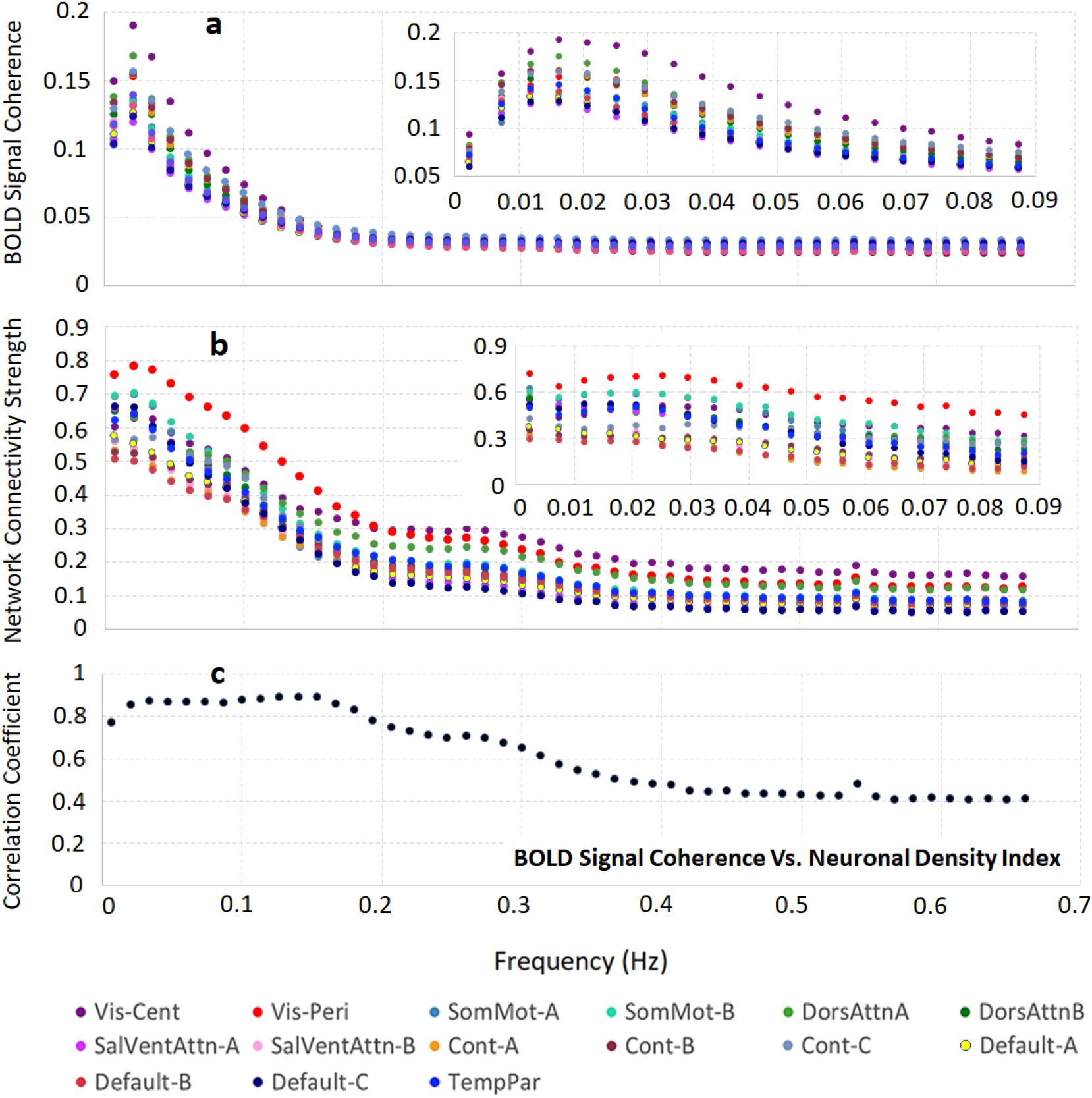
Relationship between frequency content of the resting state BOLD signal and the brain cellular structure. **(a)** BOLD signal coherence (mean standard deviations of the rs-fMRI signal) in 15 networks as a function of rs-fMRI signal frequency content. **(b)** The intra-network connectivity strength (NCS) of 15 networks as a function of rs-fMRI signal frequency content. Both panels **(a)** and **(b)** show relatively sharp peaks for all networks between 0.01 Hz and 0.03 Hz that are further illustrated in the insets. **(c)** Correlation coefficient between BOLD signal coherence in 15 networks and tissue neuronal content of these networks (as in **Figure 3**) as a function of rs-fMRI signal frequency content. The frequency resolution in main graphs is 0.014 Hz and in the inset it is 0.0047 Hz. Data show the strongest correlation in a rather broad infra-slow frequency range between 0.01 – 0.16 Hz.

## DISCUSSION

While numerous specific mechanisms involved in neuronal signal transmission between different parts of the CNS are well understood, understanding the interrelationships between these mechanisms, and how they contribute to brain function, is still an elusive goal of neuroscience. A great number of papers are now devoted to one branch of this goal - studying brain functional networks reflecting intrinsic brain activity in the resting state (e.g. see (Raichle 2015) and references therein). While significant progress has been achieved in studying resting state functional networks in health and disease, many questions related to the relationship between resting state functional organization and underlying brain cellular organization are still not well understood.

Substantial advances in this direction have been achieved by utilizing information on brain genetic organization available from the Allen Human Brain Atlas (https://portal.brain-map.org/). Richiardi et al. (Richiardi, Altmann et al. 2015) found that functional networks are underpinned by the networks of genes coding for ion channels and synaptic functions. Hawrylycz et al. (Hawrylycz, Miller et al. 2015) demonstrated that genes in the neuron-associated networks showed higher preservation between human brains and were related to functionally relevant circuitry. Furthermore, Goyal et al. (Goyal, Hawrylycz et al. 2014) were able to differentiate between the use of aerobic glycolysis in networks, which was associated with the presence of genes related to synapse formation and growth (i.e., transcriptional neotony).

By using information on spatial distribution of gene expression profiles in human brain provided by the Allen Human Brain Atlas, Wen et al. (Wen, Goyal et al. 2018) identified three *gene structural networks* related to brain neuronal, glia, and synaptic structures. Importantly, a strong association between these networks and the R2t* metric of the qGRE MRI signal (Ulrich and Yablonskiy 2016) was established in (Wen, Goyal et al. 2018). This showed that *in vivo* measurement of the major baseline tissue-cellular-specific component of qGRE signal decay rate parameter R2t* (t stands for tissue) provides a unique genetic perspective into the cellular constituents of the human cortex and can serve as a previously unidentified link between cortical tissue cellular composition and MRI signal. Analysis of the genetically-derived brain cellular composition (Wen, Goyal et al. 2018) was in good agreement with direct histological measurements by Herculano-Houzel (Herculano-Houzel 2014) and other previous findings (Elston, Benavides-Piccione et al. 2001, Collins, Airey et al. 2010, Glasser and Van Essen 2011).

It is worth noting that the method used in (Wen, Goyal et al. 2018) to establish a relationship between R2t* metric and brain tissue neuronal density is not as different from traditional histology as one might think. Indeed, in traditional histology, an MRI metric (R2t* in our case) would be correlated with the count of a histological markers specific to mature neurons and other markers specific to other cells such as astrocytes, and to structural components such as myelin. In (Wen, Goyal et al. 2018), instead of using histological markers, published gene expression profiles (<u>Microarray Data:: Allen Brain Atlas: Human Brain</u> (brain-map.org)) were used with their relationship to neurons established using available tools such as ToppGene portal (https://toppgene.cchmc.org) which includes 19061 genes in the “Cellular Component” category and 23956 genes in the “Coexpression Atlas” category. The sensitivity of the gene enrichment analysis was demonstrated using DAVID Bioinformatics Resources (https://david.ncifcrf.gov/). Results showed that gene expression profiles affiliated with neuronal cellular and subcellular compartments correlated strongly with R2t*.

In this paper we use quantitative relationships established in (Wen, Goyal et al. 2018) between the R2t* metric of the quantitative Gradient Recalled Echo (qGRE) MRI signal and the human brain cellular composition to study *in vivo* interrelationships between human brain resting state functional connectivity which is known to provide insights into large-scale brain circuit organization (Fox and Raichle 2007, Buckner, Krienen et al. 2013) and underlying brain cellular organization.

Our functional connectivity analysis is based on the resting state data obtained from the HCP1200 dataset (February, 2017 release) of young adults aged between 22 and 35 (Van Essen, Smith et al. 2013). To analyze resting state functional connectivity data we use brain parcellation into individual functional units and brain resting state networks proposed in (Yeo, Krienen et al. 2011, Schaefer, Kong et al. 2018) that allow significant reduction of large volume of resting state functional connectivity data but also providing a sufficient number of parcels (ROIs) to delineate the importance of brain cortical anatomical structures as emphasized by Van Essen et al. (Van Essen, Glasser et al. 2012).

Our results elucidate the relationship between the components of human brain cellular composition (neurons, glia cells and synapses) and the BOLD signal characteristics of the resting state functional networks. In our approach, the functional connectivity inside the individual functional units (ROIs) is characterized by the coherence of the BOLD signals from the voxels/vertices comprising functional units (Eq. [3]), while the intra-network functional connections are characterized by the coherence of the global BOLD signals generated by functional units comprising these networks (Eqs. [8], [9]). Our analysis reveals the relationships between brain functional properties and cellular content. The general trend indicates that the functional units with higher concentration of neurons and correspondingly lower concentration of glial cells and synapses display stronger coherence of the rs-fMRI BOLD signals in the individual functional units (**Figure 3** and **Figure A3 in the Appendix**). This result reveals that the synchrony of the connections between brain cellular circuits in individual functional units is mostly governed by the neurons – a higher concentration of neurons leads to stronger functional connections. At the same time, the synchrony of connections between cellular circuits belonging to different functional units in the network is largely governed, not only by the strength of the neuron-neuron associations between functional units, but also neuron-synapse, and neuron-glia associations (**Figure 4**).

The strong role of the neuronal-glia association in forming strong network connections is also in agreement with the role that glial cells play by providing metabolic (Suzuki, Stern et al. 2011, Magistretti and Allaman 2015) and regulatory (Poskanzer and Yuste 2011) support for neurons as well as support for neurite outgrowth and neuronal guidance (Ullian, Sapperstein et al. 2001, Pannasch, Vargova et al. 2011). Moreover, neuron-glia cross-talk leads to synaptic formation and remodeling. In absence of glia, neurons generate weak synapses (Araque and Navarrete 2010, Pannasch, Vargova et al. 2011). The dependence of intra-network connectivity strength on the neuron-synapse relationship could also have been expected because they operate as a unit in conducting brain electric currents.

To further clarify the role that different parts of the neuron play in formation of brain functional connectivity, we can compare our R2t*-based results with a T1w/T2w-derived proxy related to the cortical tissue myelin content proposed (Glasser and Van Essen 2011) and successfully used (Glasser, Coalson et al. 2016) for mapping human cortical areas by Glasser and Van Essen. An association between T1w/T2w myelin proxy and gene profiles from an Allen Human Brain Atlas was also established in (Burt, Demirtas et al. 2018). Data in **Figure 5a** shows a very strong (R^2^ = 0.61) correlation between measurements of the Neuronal Density Index (NDI) and T1w/T2w-derived Myelin Index (MI) across the 15 networks (Limbic networks were omitted from this consideration for the same reasons that R2t* and functional connectivity data were omitted). This result is consistent with previous findings that the neuronal densities tend to be higher in areas of high myelin content and low in areas of low myelin content (Collins, Airey et al. 2010, Glasser and Van Essen 2011) that have more complex intracortical circuitry (i.e. larger dendritic field sizes and larger dendritic arbors) (Elston, Benavides-Piccione et al. 2001). At the same time, data suggest that not all parts of the R2t*-based calculated NDI are associated with the presence of myelinated neuronal processes. These results are expected because *myelin covers only part of the neuron*, i.e. myelinated axons, while R2t*-based *NDI comprises all parts of neuronal structure*, i.e., cell body, myelinated and not-myelinated axons, and dendrites.

The conclusion that the qGRE-derived NDI also has additional contributions of myelinated axons from remaining neuronal components – dendrites and soma – is further supported by data in **Figure 5b** and **Figure 5c**. Indeed, data in **Figure 5b** show that the correlation between BOLD signal coherence and Myelin Index is much weaker (R^2^ = 0.23) than the correlation between BOLD signal coherence and NDI (R^2^ = 0.51, **Figure 3**). Subsequently, comparison of data in **Figure 4** and **Figure 5c** shows that the intra-network connectivity strength has stronger association with neuron-neuron associations (R^2^ = 0.53) than with the associations due to the myelinated processes (R^2^ = 0.47). This might suggest that the qGRE NDI metric has a stronger association with the brain cellular “functionality” than the T1w/T2w-derived myelin index. These results are quite instructive. They suggest that the BOLD signal coherence in the individual brain ROIs comprising functional connectivity networks might be mostly associated with the non-myelinated parts of the neurons – dendrites and soma. At the same time, the inter-ROIs connectivity has significantly stronger contribution from myelinated axons needed to secure inter-unit, within-network functional connectivity.

In this paper we have also provided a detailed analysis of the “frequency content” of the resting state BOLD signal and a corresponding strength of the resting state functional connectivity in relation to the underlying cellular composition. It is well appreciated that the resting state BOLD signal originates from a specific element of brain activity, i.e., *infra-slow* activity with frequencies approximately below 0.1 Hz which is thought to be linked in this interval to the neuronal activities (Fox and Raichle 2007, Zhang and Raichle 2010, Han, Wang et al. 2011, Palva and Palva 2012, Mitra, Kraft et al. 2018, Raut, Snyder et al. 2020). Our data are in agreement with this notion – for all networks, the resting state BOLD signal coherence and the intra-network connectivity strength are not monotonic functions of the frequency with a rather sharp peaks in the *infra-slow frequency* range between 0.01 and 0.16 Hz (**Figure 6a** and **Figure 6b**). A relationship of these infra-slow BOLD signal fluctuations to neuronal activity is clearly supported by a plot in **Figure 6c** showing a high correlation plateau in the frequency range of 0.01 to 0.16 Hz on the curve displaying correlation between Neuronal Density Index and BOLD signal coherence.

All these mechanisms lead to a rather broad distribution of resting state functional networks’ properties. We found that the visual networks express the highest neuronal density, yet lowest density of glial cells and synapses, and the strongest intra-functional-unit BOLD signal coherence and intra-network connectivity. The DMN shows relatively low intra-functional-unit BOLD signal coherence and intra-network connectivity strength, but significant diversity of cellular constituents. The DMN part B exhibits a remarkably balanced cellular content – Neuronal Density Index (0.51) was almost exactly equal to Glia Density (0.49) and Synaptic Density (0.48) indices. At the same time, the other parts of DMN (A and C) that include temporal and inferior parietal lobules, dorsal prefrontal, precuneus posterior cingulate, medial prefrontal, retrosplenial, and parahippocampal cortices have more condensed neuronal content with relatively lower concentration of synapses and glia cells. This network is affiliated with temporal, inferior parietal, dorsal prefrontal, lateral prefrontal, and ventral prefrontal brain regions. These results can potentially be helpful in understanding the unique features of DMN and its very prominent role in the overall organization of the brain (Margulies, Ghosh et al. 2016) plus it’s the target of Alzheimer disease (Palmqvist, Schöll et al. 2017).

As the study of resting state functional connectivity within and among the large networks of the human brain has moved forward, the concept of hierarchies among these networks has emerged as a major area of interest (for example see (Huntenburg, Bazin et al. 2018, Raut, Snyder et al. 2020) and the references therein). Broadly speaking, networks are positioned according to a variety of features beginning with the sensorimotor cortices and ascending to association cortices with the brain’s default mode network at the top of this hierarchy. Networks at the top of the hierarchy are less myelinated (Glasser, Goyal et al. 2014), exhibit higher levels of aerobic glycolysis (Vaishnavi, Vlassenko et al. 2010, Glasser, Goyal et al. 2013) and harbor neotenous genes (Goyal, Hawrylycz et al. 2014). Together these features relate to the presence of synapses and their role in growth and plasticity. Our data would add that networks at the top of the hierarchy have denser synaptic relative to neuronal structure (see **Figure 2**). This may be relevant to our understanding of a functional feature of networks at the top of the hierarchy as they integrate incoming information more slowly than, for example, primary sensory areas (Chaudhuri, Knoblauch et al. 2015, Raut, Snyder et al. 2020).

## CONCLUSION

In this paper we have provided a detailed analysis of the associations between human brain cellular composition and functional networks identified by the resting state BOLD signal. Our results show that the brain regions with higher concentration of neurons, but relatively lower concentration of glial cells and synapses support strong connections between cellular circuits in the network-defined functional units, leading to the strong BOLD signal coherence in individual functional units. Furthermore, our results show that a significant contribution to a between-unit connectivity is provided by the neuron-neuron, neuron-synaptic, and neuron-glia associations between cellular circuits.

These relationships lead to a rather broad distribution of the resting state functional networks’ properties. We found that the visual networks have the strongest BOLD signal coherence with the highest neuronal density (but lowest density of glial cells and synapses), and the strongest intra-network connectivity. The Default Mode Network (DMN) shows relatively low BOLD signal coherence and Network Connectivity Strength but significant diversity of cellular circuits reflecting the DMN’s prominent role in the overall organization of the brain and the hierarchy of functional networks.

## AUTHOR CONTRIBUTIONS

S.K., M.E.R., and D.A.Y. designed research; S.K., M.E.R. and D.A.Y. performed research; S.K. and D.A.Y. analyzed data; and S.K. and D.A.Y. wrote the paper.

## ACKNOWLEDGEMENTS

The resting state data of this study were provided by the Human Connectome Project, WU-Minn Consortium (Principal Investigators: David Van Essen and Kamil Ugurbil; 1U54MH091657) funded by the 16 NIH Institutes and Centers that support the NIH Blueprint for Neuroscience Research; and by the McDonnell Center for Systems Neuroscience at Washington University. The genetic information was obtained from Allen Institute for Brain Science, Allen Human Brain Atlas, Available from: human.brain-map.org. The authors would also like to acknowledge helpful discussions at different stages of this project with Charles Hildebolt, Dillan Newbold, Reza Hamidian, Janine Bijsterbosch, BT Thomas Yeo and Ryan Raut. Editing assistance was provided by InPrint: A Scientific Editing Network at Washington University in St. Louis.

## APPENDIX

**Figure A1.**
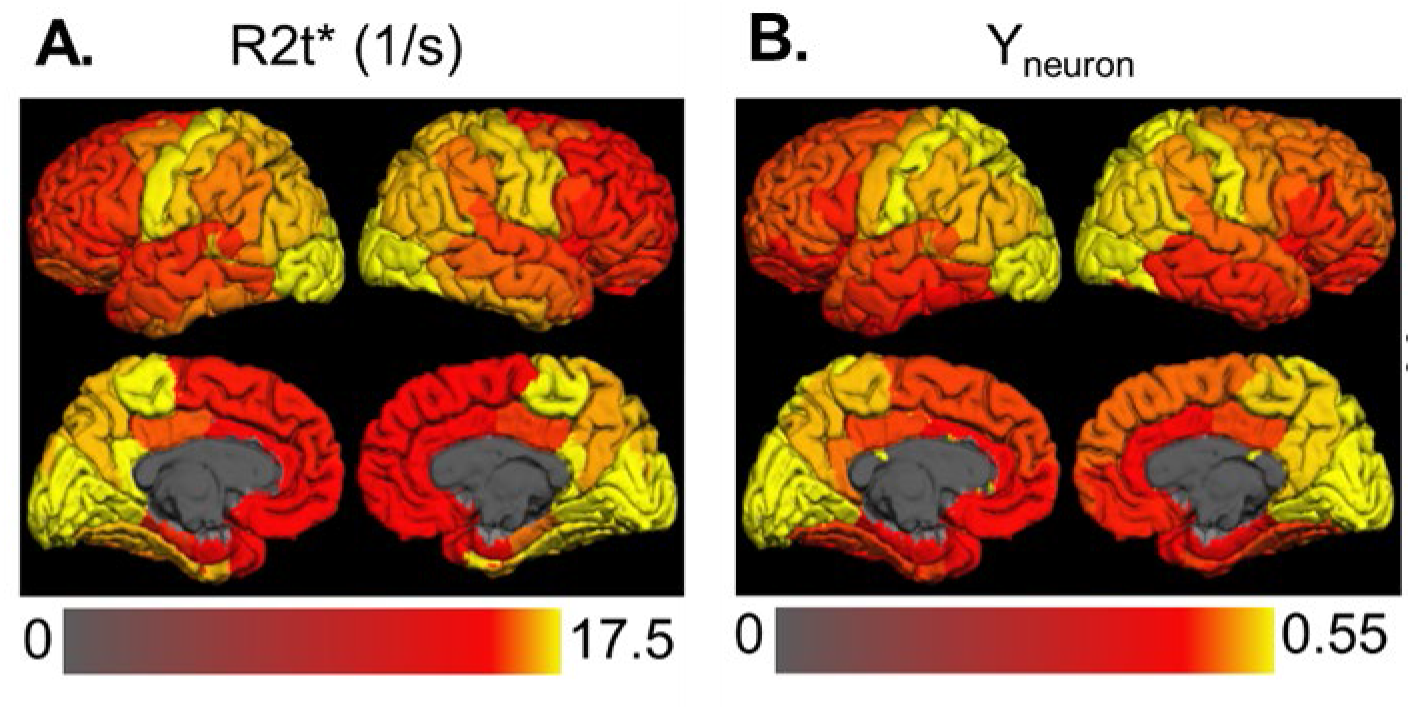
The association between the expression of neuron-related genetic network across different regions of the human brain and R2t* measurements in the same regions (adapted from (Wen, Goyal et al. 2018)). R2t* map (**A**) in this figure is obtained from qGRE MRI; Yneuron (**B**) (designated as Neuronal Density Index) is obtained from the analysis of the gene expression profiles in the Allen Human Brain Atlas – two totally independent measurements. The figure and analysis in (Wen, Goyal et al. 2018) provided convincing and consistent with literature arguments that the R2t* metric of qGRE signal is strongly associated with the neuronal density.

**Figure A2.**
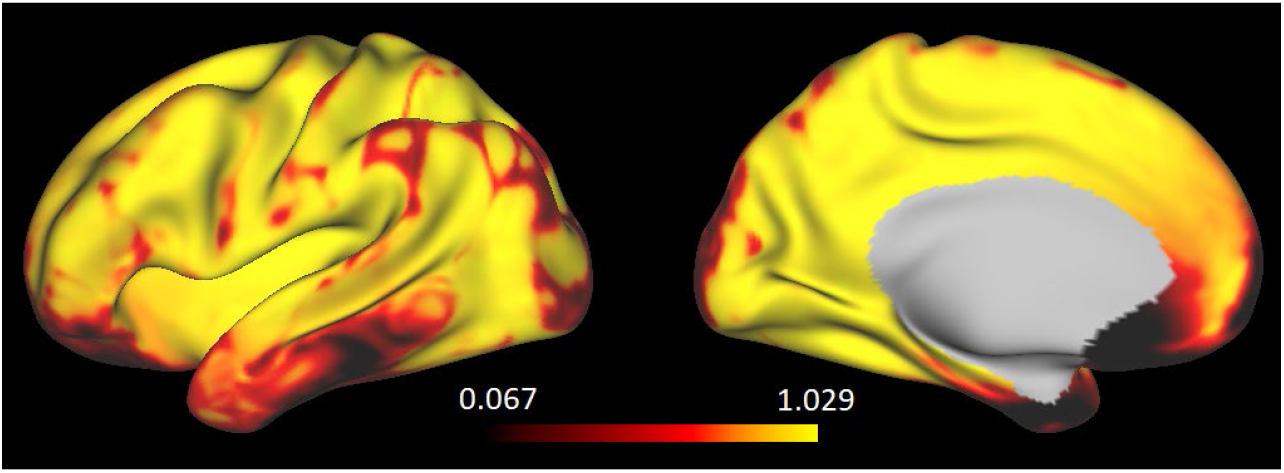
Map characterizing magnetic field inhomogeneities in the human brain. Data obtained from the phase information of qGRE signal by means of the Voxel Spread Function approach (Yablonskiy, Sukstanskii et al. 2013). Yellow color designates areas lightly affected by magnetic field inhomogeneities. Black color designates severely affected areas. The scale bar shows values of the so-called F-function (Yablonskiy, Sukstanskii et al. 2013) at the gradient echo time TE =40 ms. F-function shows fraction of signal not affected by the loss due to the field inhomogeneities, ranging from 1 (no signal loss) to 0 (full loss of signal). Images show significant loss of signal in both Limbic networks, thus adversely affecting quantitative analysis of both, rs-fMRI and qGRE data.

**Figure A3.**
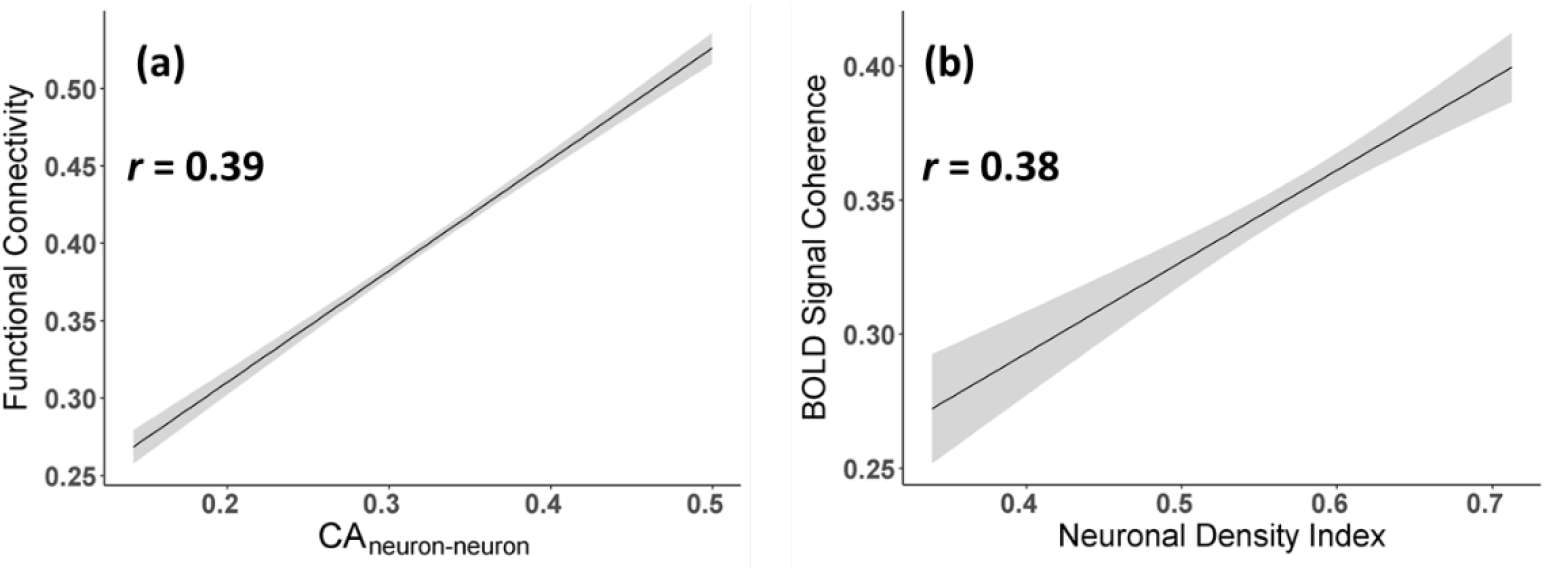
Functional-Structural Associations. **(a):** Association between Functional Connectivity Strength defined based on 280 ROIs and cellular association between neurons in these ROIs. The correlation is highly significant with p= 9 E-98; **(b):** Association between BOLD signal coherence in 280 ROIs and neural density index in these ROIs. The correlation is highly significant with p= 5.3 E-11. In both curves ROIs in limbic networks are omitted. Shaded areas – 95% confidence intervals. Actual data are not shown due to their big number.

**Figure A4.**
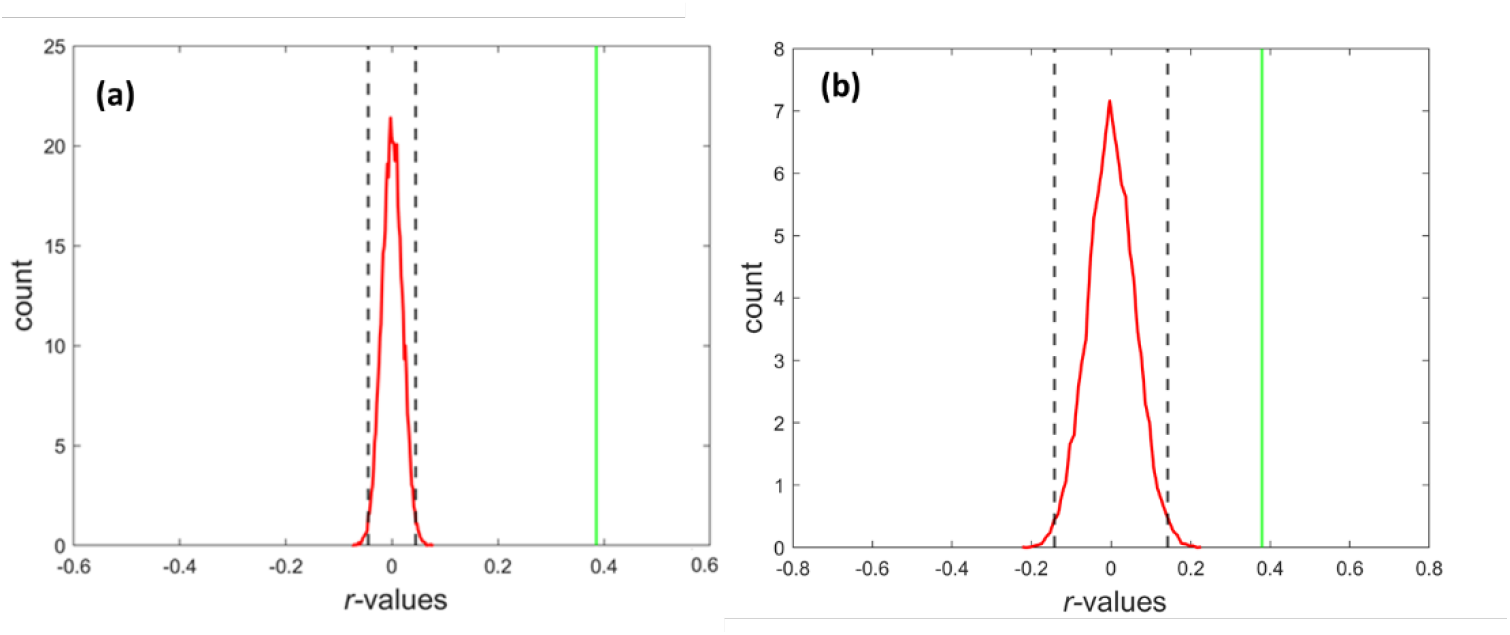
Permutation Test Analysis (randomization test). **(a)**: Functional Connectivity vs. Structural Connectivity between neurons, Y (neuron-neuron). The curve shows the distribution of *r*-values calculated between Y (neuron-neuron) and randomly permuting FC 20000 times. **(b)**: BOLD Signal Coherence vs. Neuronal Density Index (NDI). The curve shows the distribution of *r*-values calculated between NDI and randomly permuting 10000 times BOLD Signal Coherence. In both plots, the 99% percentiles of the distribution lines are marked by black dashed lines, the r-values without permutation are marked as green lines and are well outside black lines. The result shown in the figures validates a significant difference in both cases with p-values <0.000001).

**Figure A5.**
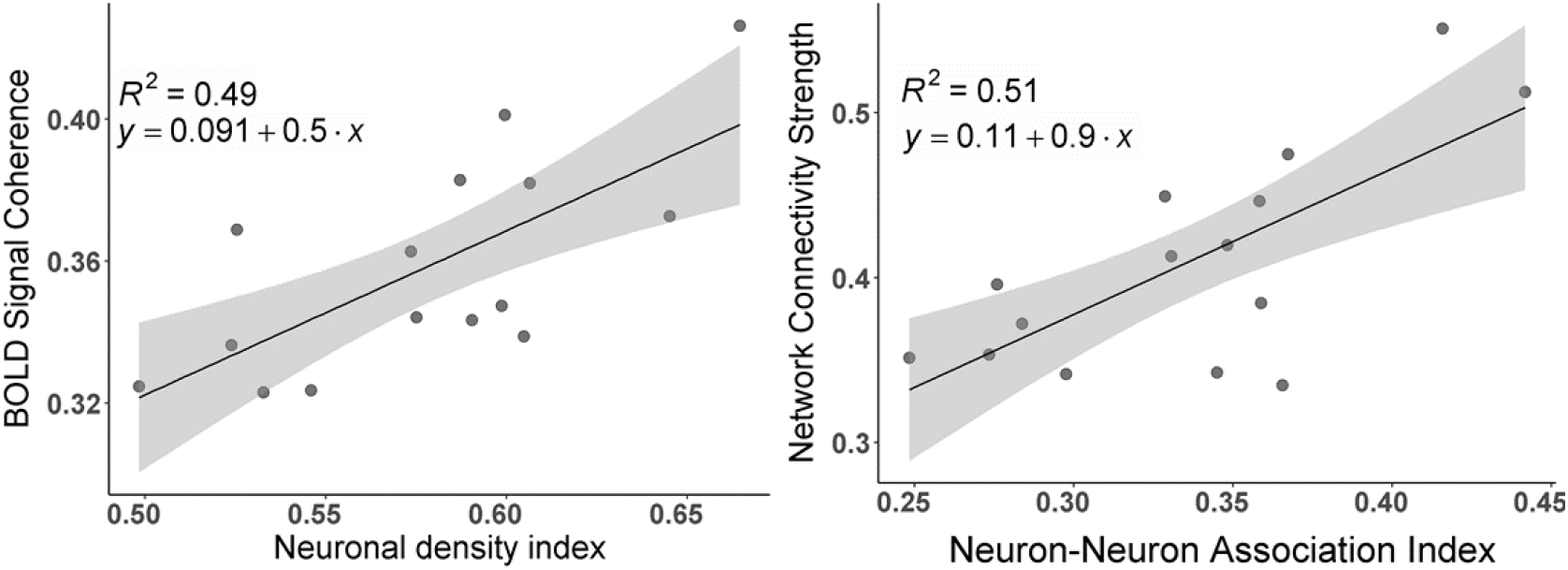
Statistical sufficiency of sample size. To demonstrate statistical sufficiency of our sample size (N=183) we have run analysis similar to presented in **Figure 3** and **Figure 4** of the main text using a smaller sample size (N = 100). Results presented herein in **Figure A5** show practically the same correlations with only slightly smaller R^2^ values.

